# Stimulation of the muscarinic receptor M4 activates quiescent neural precursor cells and ameliorates medial septum cholinergic lesion-induced impairments in adult hippocampal neurogenesis

**DOI:** 10.1101/2022.08.25.505357

**Authors:** Lidia I. Madrid, Saurabh Bandhavkar, Katelyn Hafey, Javier Jimenez-Martin, Michael Milne, Elizabeth J. Coulson, Dhanisha J. Jhaveri

**Author notes:** **Corresponding authors:** Dr Dhanisha J. Jhaveri -, Prof. Elizabeth Coulson.

## Abstract

Cholinergic signaling plays a crucial role in the regulation of adult hippocampal neurogenesis and hippocampus-dependent cognitive and mood-related functions. However, the contribution of basal forebrain medial septum (MS) and diagonal band of Broca (DBB) cholinergic neurons that innervate the hippocampus and the identity of the cholinergic receptor(s) that regulate the production and maturation of new neurons are not completely understood. Using a targeted, selective ablation approach, we show that MS/DBB cholinergic neurons support both the survival and morphological maturation of adult-born neurons in the mouse hippocampus. We demonstrate that the muscarinic acetycholine receptor subtype M4 (M4 mAChR) is expressed on a population of quiescent neural precursor cells (NPCs) and that its pharmacological stimulation via intra-hippocampal or systemic administration of M4-selective modulators leads to their activation, thereby enhancing neurogenesis *in vivo*. Furthermore, we show that the activation of M4 mAChR-expressing quiescent NPCs ameliorates the MS/DBB cholinergic lesion-induced decrease in hippocampal neurogenesis. In contrast, the impairment in the morphological maturation of adult-born neurons due to MS/DBB cholinergic neuron loss is further exacerbated by the systemic administration of an M4-selective allosteric potentiator. These findings reveal novel and stage-specific roles of cholinergic signaling in regulating adult hippocampal neurogenesis. They also uncouple the positive role of selective M4 potentiators in enhancing the production of new neurons from the M4-induced inhibition of their morphological maturation, at least in the context of cholinergic dysfunction.

**Significance statement:** Cholinergic signaling plays an important role in the regulation of adult hippocampal neurogenesis and cognitive function, with impairments in these processes reported as early pathogenic events in age-related dementia. Here, we uncover the presence of cholinergic-responsive hippocampal precursor cells that are are directly activated by selective stimulation of the muscarinic receptor subtype M4. Furthermore, M4-mediated stimulation rescues the decrease in the level of hippocampal neurogenesis following the sepal-hippocampal cholinergic neuron loss. We also reveal that septal-hippocampal cholinergic dysfunction impairs the structural maturation of hippocampal adult-born neurons, an effect which is further exacerbated by M4 receptor modulators. These findings reveal stage-specific roles of cholinergic signaling in regulating functionally relevant adult hippocampal neurogenesis.

## Introduction

The hippocampus is a key brain region in which the developmental program of neurogenesis remains functional during adult life (Ming and Song, 2011). The heightened plasticity of these adult-born neurons has been shown to impact neuronal activity within the local circuitry and contribute to the regulation of select cognitive and mood-related functions (Christian et al., 2014). The process of adult hippocampal neurogenesis, which encompasses the proliferation of resident neural precursor cells (NPCs), their differentiation into neurons, and their integration into the existing circuitry, is sensitive to neural activity-mediated regulation (Ming and Song, 2011; Jhaveri, 2012). Amongst a range of intrinsic and extrinsic factors, several neurotransmitters, including glutamate, γ-aminobutyric acid (GABA), serotonin, norepinephrine and dopamine, have been shown to regulate various stages of adult neurogenesis, serving both non-synaptic trophic and synaptic roles (Jhaveri et al., 2010; Song et al., 2012; Berg et al., 2013; Jhaveri et al., 2015).

The neurotransmitter acetylcholine has also been linked to the regulation of adult hippocampal neurogenesis, as well as hippocampus-dependent cognitive processes (Cooper-Kuhn et al., 2004; Mohapel et al., 2005; Madrid et al., 2021). The dentate gyrus, which harbors populations of quiescent and active NPCs (Walker et al., 2008; Lugert et al., 2010; Jhaveri et al., 2015), is richly innervated by cholinergic fibers that originate from the cell bodies residing in the medial septum (MS) and the diagonal band of Broca (DBB) nuclei of the basal forebrain cholinergic system (Kaneko et al., 2006). Notably, retrograde tracing experiments in animals have shown that, during their immature phase, adult-born hippocampal neurons receive direct synaptic innervation from the MS cholinergic neurons (Vivar et al., 2012; Deshpande et al., 2013).

Given that the loss of basal forebrain cholinergic neurons, including those which innervate the hippocampus (Kerbler et al., 2013; Schmitz and Nathan Spreng, 2016), and a progressive decline in hippocampal neurogenesis (Moreno-Jimenez et al., 2019) are amongst the early pathogenic events that correlate with the cognitive impairments observed in Alzheimer’s disease, a number of studies have explored the link between cholinergic signaling and adult hippocampal neurogenesis in animal models. Supporting the neurogenic role of septal cholinergic innervation, global lesion of the basal forebrain cholinergic neurons in rats has been shown to impair hippocampal neurogenesis by reducing the survival of newborn neurons (Cooper-Kuhn et al., 2004; Mohapel et al., 2005). Corroborating these findings, the inhibition of cholinergic activity using the muscarinic antagonist scopolamine was found to decrease cell proliferation in the adult hippocampus (Chen et al., 2018), whereas intracerebroventricular infusions of oxotremorine, a selective muscarinic agonist, led to an increase in proliferation (Cooper-Kuhn et al., 2004; Mohapel et al., 2005; Van Kampen and Eckman, 2010). Interestingly, the acetylcholinesterase inhibitor donepezil, which is clinically used to treat cognitive deficits in Alzheimer’s disease, has been shown to enhance hippocampal neurogenesis (Kotani et al., 2006).

Although global basal forebrain cholinergic lesions and pharmacological approaches have suggested an important role of cholinergic signaling in regulating adult hippocampal neurogenesis (Cooper-Kuhn et al., 2004; Mohapel et al., 2005), several outstanding questions remain. Here, we asked what contributions the direct cholinergic MS/DBB innervation to the hippocampus makes to the regulation of various stages of adult neurogenesis in the mouse, and whether quiescent NPCs in the hippocampus express cholinergic receptors such that their activity can be exploited to enhance neurogenesis, including under conditions where there is cholinergic neuron loss or dysfunction.

## Materials and methods

### Animals

Adult (7-9 weeks old) male C57Bl/6J mice (Animal Resources Centre, Australia) were used to examine the effects of selective MS/DBB cholinergic lesion and pharmacological manipulation of the acetycholine receptors on adult hippocampal neurogenesis *in vitro* and *in vivo*. Male and female Nestin-GFP mice were used for fluorescence-activated cell sorting (FACS), and RNA-seq transcriptome analysis. These mice express green fluorescent protein (GFP) under the control of the Nestin promoter, which allows visualization and purification of resident NPCs (Jhaveri et al., 2015). Tamoxifen-inducible mice under the control of Achaete-scute complex homolog 1 (Ascl1)-Cre^ERT2^ (Ascl1-Cre^ERT2^; Jackson Laboratory, #012882) crossed with CAG floxStop-tdTomato reporter mice (tdTom; obtained from the Jackson Laboratory, #007914) were used to examine the morphological maturation of adult-born neurons. Mice were housed with up to 4 same-sex littermates in individually ventilated cages. All cages were given bedding and nesting material. Mice were maintained on a 12 h light-dark cycle (lights on at 07:00 h) and supplied with *ad libitum* access to water and food. Experiments were performed in accordance with the Australian Code of Practice for the Care and Use of Animals for Scientific Purposes and were approved by the University of Queensland Animal Ethics Committee (MRI-UQ/TRI/163/17, QBI/566/18).

### Medial septum cholinergic lesion

7-9 week old C57Bl/6J and Ascl1-Cre^ERT2^::tdTom mice were anesthetized with an intraperitoneal (i.p.) injection of ketamine/xylazine (50mg/kg and 8mg/kg body weight, respectively). Infusion of murine-p75 neurotrophin receptor (p75NTR)-Saporin (hereafter abbreviated as p75-Sap; 0.4 µg/µl; Advanced Targeting System) or control rabbit-IgG-Saporin (IgG-Sap; 0.4 µg/µl) was performed using a 30G needle attached to a 5 µl Hamilton syringe. The stereotaxic coordinates from Bregma for MS injections were anterior/posterior: −0.9 mm, medial/lateral: 0 mm and dorsal/ventral: −4.7 mm. The toxin was infused at a rate of 0.4 μl/min (the total volume injected was 1.5 μl), with the needle left in place for 10 min to allow for diffusion. Animals were administered Temgesic (Buprenorphine; 0.l mg/kg; subcutaneous injection; Temvet, Ilium) for postoperative pain relief.

### Labeling newborn cells

To genetically label NPCs and trace their progeny, Ascl1-Cre^ERT2^::tdTom mice were administered tamoxifen (Sigma-Aldrich) i.p. at 150 mg/kg for two consecutive days. The tamoxifen was dissolved in corn oil (Sigma-Aldrich) with 10% ethanol at 50 mg/ml concentration. To label proliferating cells *in vivo*, 5-bromo-2-deoxyuridine (BrdU; 100 mg/kg; i.p., Sigma-Aldrich) was injected daily for five consecutive days.

### Intrahippocampal and systemic stimulation of M4 mAChRs

To selectively stimulate hippocampal muscarinic acetycholine receptor subtype M4 (M4 mAChRs), 10 μM VU10010 (Tocris), a selective allosteric potentiator (Shirey et al., 2008) was directly infused into the hippocampus using a single cannula attached to a 7 day, 0.5 μl/h osmotic pump (Alzet Model 1007D). Vehicle (0.1% DMSO, 0.2% bovine serum albumin in saline) was infused in the control group of mice. The coordinates from Bregma were: anterior/posterior: −2.0 mm, medial/lateral: −1.3 mm and dorsal/ventral: −2.1 mm. The osmotic pump was subcutaneously implanted caudal to the scapula. The mice were perfused 15 days after the end of the infusion period and their brains were harvested for immunohistochemical analysis. To evaluate the effects of the blood-brain barrier-permeable, potent and selective Chrm4 allosteric modulator VU0152100 (Brady et al., 2008) on adult hippocampal neurogenesis, mice were treated with VU0152100 (0.5 mg/kg, Sigma-Aldrich) or vehicle via daily i.p. injections for 7 consecutive days. The VU0152100 was dissolved in 40% PEG800, 5% Tween20, 10% DMSO in saline. Mice were perfused 15 days after the end of the treatment and their brains were collected for histology.

### Tissue collection and immunohistochemistry

Mice were perfused transcardially using ice-cold 4% paraformaldehyde. Their brains were removed and post-fixed in 4% paraformaldehyde for 2 h, then left in 30% sucrose for up to 48 h, after which 40 μm sections were cut using a freezing microtome. The sections were washed in 0.1M PBS three times and stored at 4 °C with 0.05% sodium azide. Every sixth section was used for immunohistochemical analysis. For BrdU immunolabeling, the sections were rinsed in 0.1M PBS and incubated in 1M HCl at 45 °C for 20 min. Sections were rinsed in boric acid (pH 6) for 5 min, followed by a wash in 0.1M PBS. They were then placed into a blocking solution containing 0.1% Triton X-100 in PBS (0.1% PBST) and 5% normal goat serum for 1.5 h at room temperature. The sections were subsequently incubated overnight in solution containing primary antibodies, including mouse anti-BrdU antibody (1:500; Roche), rabbit anti-p75NTR (1:1000; Millipore 07-476), rabbit anti-doublecortin (DCX; 1:500; Cell Signalling Technology), rabbit anti-Ki67 (1:500; Novacastra), goat anti-tdTomato (Accurate Chemical and Scientific Corporation; 1:500,) and DAPI (1:5000; Life Technologies). The sections were then washed in 0.1% PBST and incubated in the appropriate species-specific secondary antibody for 2 h at room temperature (goat anti-rabbit Alexa 488, 1:2000; goat anti-mouse Alexa 647, 1:2000; donkey anti-goat Alexa 568; 1:2000) and 4′,6-diamidino-2-phenylindole (DAPI, 1:1,000, Life Technologies). Following several washes, the sections were mounted using fluoromount (DakoCytomation) and viewed on a Zeiss-Axio Imager microscope and Diskovery spinning disk confocal microscope (Andor Technology) built around a Nikon Ti-E body (Nikon Corporation) and equipped with two Zyla 4.2 sCMOS cameras (Andor Technology) and controlled by Nikon NIS software.

### Stereology and quantification

An upright stereology microscope (MicroBrightField Bioscience) built around a Zeiss Axio Imager.Z2 upright microscope body with an ORCA-R2 digital charge-coupled device camera (Hamamatsu Photonics K.K.) for fluorescence imaging and a 20x objective lens (Zeiss Plan-ApoChromat 20x/0.8 NA air objective) was used to perform cell counts. Stereo Investigator software (version 2017.03.3; MicroBrightField Bioscience) was used to quantify cells expressing single markers (p75NTR, BrdU, DCX, or Ki67) as well as those showing co-localization (BrdU and DCX). A minimum of six sections per brain were used and cell quantification was performed in a blinded manner with regards to experimental conditions. Cell numbers in the dentate gyrus were divided by the length of the subgranular zone and the data were expressed as ‘percentage change’ relative to the control.

### Microscopy and image analysis

To visualize p75NTR^+^ cells in the basal forebrain nuclei, images were acquired using an upright fluorescence slide scanner (Metafer VSlide Scanner by MetaSystems using Zeiss Axio Imager Z2) with a 20x air objective. p75NTR^+^ cells in the MS/DBB, nucleus basalis of Meynert (NBM) and substantia innominata (SI) were quantified using Imaris software (Version 9.3). For experiments evaluating neurogenesis and adult-born neuron morphology, only animals with more than 50% loss of p75NTR^+^ cells in the MS/DBB following p75-Sap injection were included in the study. Fluorescence immunolabeling in tissue was visualized and imaged using a Nikon Plan Apochromat 20x/0.75 NA air objective and a Plan Apo Lambda 60x/1.4 NA oil-immersion objective on a Diskovery spinning disk confocal microscope. Representative images of BrdU, DCX, DAPI, or tdTomato staining in the dentate gyrus were taken using a 20x objective at 2 μm intervals. For morphological analysis, three-dimensional images of dendrites were obtained from Z-stacks of confocal images taken at 0.15 μm intervals using a 60x oil immersion objective. To accurately reconstruct the dendritic morphology, we selected tdTom^+^ newborn neurons that were entirely contained within the tissue section. Morphological analysis was performed using the Imaris software.

### Neurosphere assay

8-9 week-old male C57Bl/6J mice were sacrificed by cervical dislocation and their brains were removed and collected in ice-cold Hank’s essential medium. The hippocampi were microdissected and minced and a single-cell suspension was prepared as previously described (Jhaveri et al., 2015). The resulting pellet was resuspended in 1 ml of complete neurosphere medium containing epidermal growth factor (EGF, 20 ng/ml; receptor grade, BD Biosciences) and basic fibroblast growth factor (bFGF, 10 ng/ml; recombinant bovine, Roche). The cells were then plated in a 96-well plate in complete neurosphere medium containing DMEM/F-12 EGF and bFGF, with or without muscarine (10 μM, 50 μM), nicotine (10 μM, 50 μM), VU010010 (10 μM), VU0152100 (10 μM), and PD102807 (1 μM). Muscarine was dissolved in 100% DMSO, with the final concentration of DMSO being adjusted to 0.1% for each treatment, including vehicle. The plates were incubated at 37 °C and the total number of neurospheres obtained in each treatment group was determined on day 14, with all conditions normalized to the control group for each experimental replicate and plotted as a percentage of the control.

### Fluorescence-activated cell sorting

Brains from 7–9 week-old Nestin-GFP mice were used for FACS. A single-cell suspension of hippocampal tissue was prepared as described above. The resulting pellet was resuspended in DMEM/F12 medium and incubated with biotinylated EGF conjugated with Alexa Fluor 647-streptavidin (EGF-647; 2 μg/ml; Life Technologies) for 30 to 40 min at 4°C. The antibody was washed off using excess DMEM/F12 medium. Propidium iodide (1μg/ml) was then added to label and exclude dead cells. The cell suspension was filtered using a 40µm filter prior to sorting. Cells were analyzed and sorted using a FACS Aria sorter (Becton Dickinson). The positive gates were set relative to the basal fluorescence levels obtained from wild type littermates and single fluorescence controls. Nestin GFP^+^ cells co-expressing epidermal growth factor receptor (Nes-GFP^+^EGFR^+^), which represent a near pure population of NPCs (Jhaveri et al., 2015), were sorted and collected in a tube containing neurosphere medium. Cells were plated in 96-well plates at a clonal density (<1 cell per well) in complete neurosphere medium. A total of 24 wells were plated per condition (control, muscarine, muscarine + VU10010). Neurospheres were counted on day 14 and conditions were normalized to the control group.

### Statistical analysis

Statistical analyses were performed using GraphPad Prism 9 (GraphPad). Normally distributed data were analyzed using a two-tailed Student’s t-test when comparing two groups or a one-way or two-way ANOVA followed by Bonferroni’s multiple comparison post hoc test when comparing more than two groups. For non-normally distributed data, a Log-rank test was used. Differences with a *p-*value of < 0.05 were considered statistically significant. All data are presented as the mean ± SEM.

## Results

### Selective ablation of MS/DBB cholinergic neurons leads to loss of cholinergic innervation to the hippocampus

The basal forebrain cholinergic system comprises the NBM, the SI, the horizontal and vertical DBB and the MS, with the MS/DBB providing the primary source of cholinergic innervation to the hippocampus (Auld et al., 2002). Although previous studies have shown that a global basal forebrain cholinergic lesion negatively impacts adult hippocampal neurogenesis (Cooper-Kuhn et al., 2004; Mohapel et al., 2005), the precise contribution of the direct innervation from the MS/DBB to newborn neurons remains unknown. A characteristic feature of all basal forebrain cholinergic neurons is the expression of high levels of p75NTR (Woolf et al., 1989), which can be targeted using the immunotoxin murine-p75-Saporin (p75-Sap), a monoclonal antibody conjugated to the ribosome-inactivating protein saporin (Moreau et al., 2008). Intracerebroventricular injections of p75-Sap have previously been used to effectively ablate basal forebrain cholinergic neurons of all nuclei or of specific subnuclei (Hamlin et al., 2013). To selectively ablate only the MS/DBB cholinergic neurons, we injected p75-Sap or a rabbit-Saporin control (IgG-Sap) directly into the MS/DBB (Fig. 1A). Two weeks after the MS/DBB injection, we observed a profound reduction in the number of MS/DBB cholinergic neurons in animals injected with p75-Sap compared to those injected with IgG-Sap (Fig. 1B, C). In contrast, the number of cholinergic neurons in the NBM and SI remained unaltered, establishing the specificity of the lesion (Fig. 1D, E). The significant reduction in MS/DBB cholinergic neurons was accompanied by a substantial decrease in the cholinergic fiber density in both the molecular and granular cell layers of the hippocampus as well as in the hilar region (Fig. 1F-J). Thus, direct infusion of p75-Sap into the MS/DBB induces near complete and selective loss of cholinergic neurons and their innervation of the hippocampus, while preserving other basal forebrain cholinergic nuclei.

**Figure 1.**
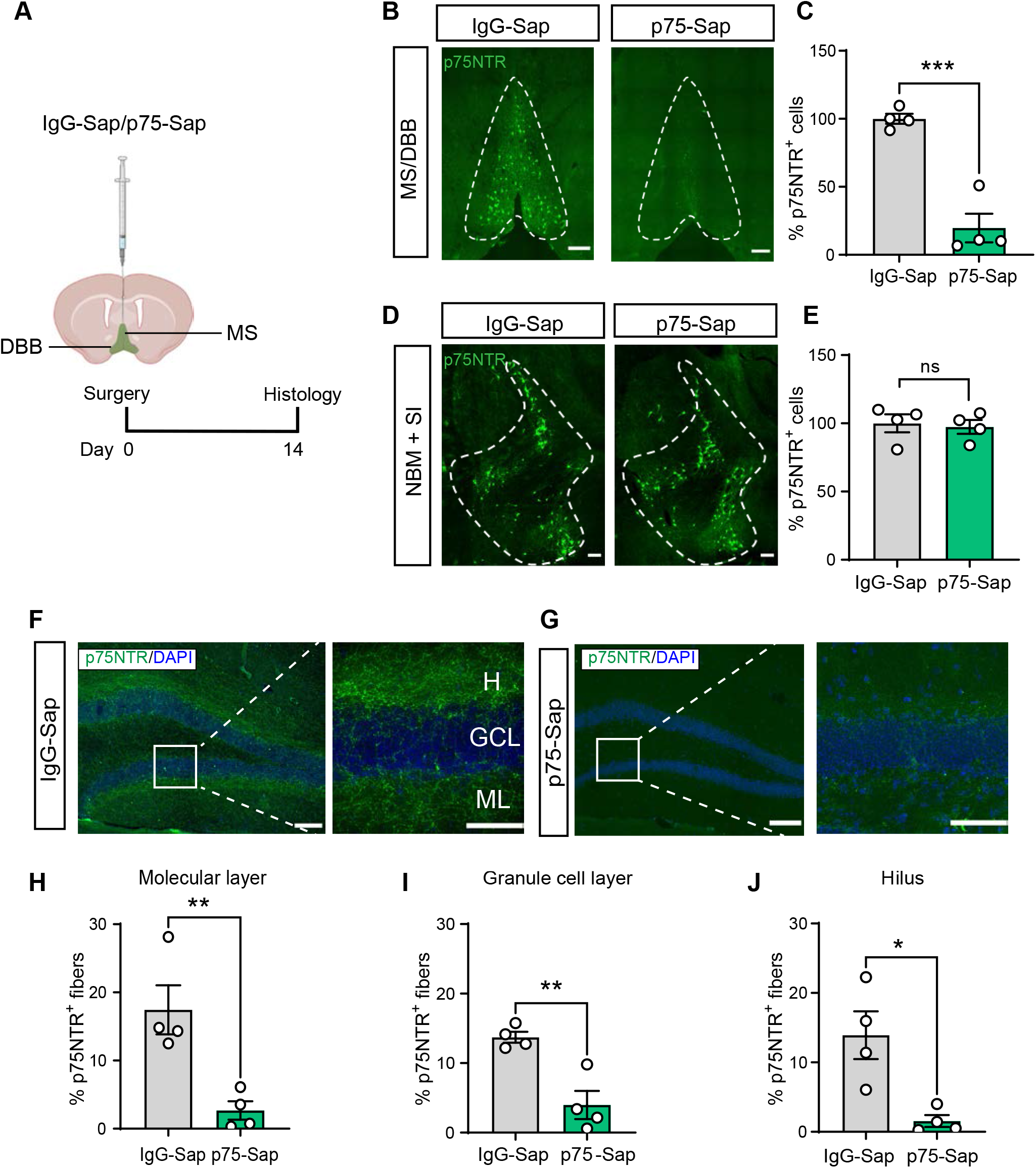
Selective ablation of MS/DBB cholinergic neurons. (A) Experimental timeline showing selective ablation of MS/DBB cholinergic neurons in the mouse brain by direct infusion of p75-Sap. Control mice were infused with IgG-Sap. (B) Representative images displaying loss of p75NTR^+^ (green) cholinergic neurons in the MS/DBB of p75-Sap-treated mice compared to control mice. (C) Quantification showing ablation of p75NTR^+^ cholinergic neurons in the MS/DBB in p75-Sap-compared to IgG-Sap-infused mice. (D) Representative images showing p75NTR^+^ neurons in the NBM and SI. (E) No loss of NBM+SI cholinergic neurons was observed in p75-Sap-compared to the IgG-Sap-treated mice. (F) A confocal image displaying p75NTR^+^ fibers (green) and nuclei stained with DAPI (blue) in the hippocampus in IgG-Sap-treated and (G) p75-Sap-treated mice. Box shows hippocampal subregions in which these fibers were quantified. H: hilus, GCL: granule cell layer, ML: molecular layer. Quantification of p75NTR^+^ fibers in the (H) molecular layer, (I) granule cell layer and (H) hilus show their profound loss in p75-Sap-infused mice. Data represent mean ± SEM. \**p* < 0.05, \*\**p* < 0.01, \*\*\**p* < 0.001; unpaired Student’s t-test. Scale bars: 100 µm.

### Selective MS/DBB cholinergic lesion reduces the survival of newborn neurons in the hippocampus

To evaluate the effects of selective MS/DBB cholinergic lesion on adult hippocampal neurogenesis, we next labeled proliferating cells using BrdU two weeks post-lesion, with the mice being sacrificed 3 weeks later (Fig. 2A). We found that the MS/DBB lesion resulted in a significant decrease in hippocampal neurogenesis, with fewer BrdU^+^ cells and newly generated neurons (BrdU^+^/DCX^+^) observed in p75-Sap-compared to IgG-Sap-injected mice (Fig. 2B-D). A significant reduction in the total number of DCX^+^ immature neurons was also observed in these mice (Fig. 2E), with no difference noted in the proportion of proliferating cells that differentiated into neurons (data not shown). We also found no difference in the number of cells expressing Ki67, an endogenous marker of cell proliferation, between groups (Fig. 2F). Together, these data indicate that the loss of cholinergic innervation into the hippocampus does not impact the baseline proliferation or neurogenic differentiation of NPCs. Rather this innervation is important for the survival of any newly generated progenitors and/or neurons, likely through direct inputs from MS/DBB cholinergic afferents (Vivar et al., 2012; Deshpande et al., 2013).

**Figure 2.**
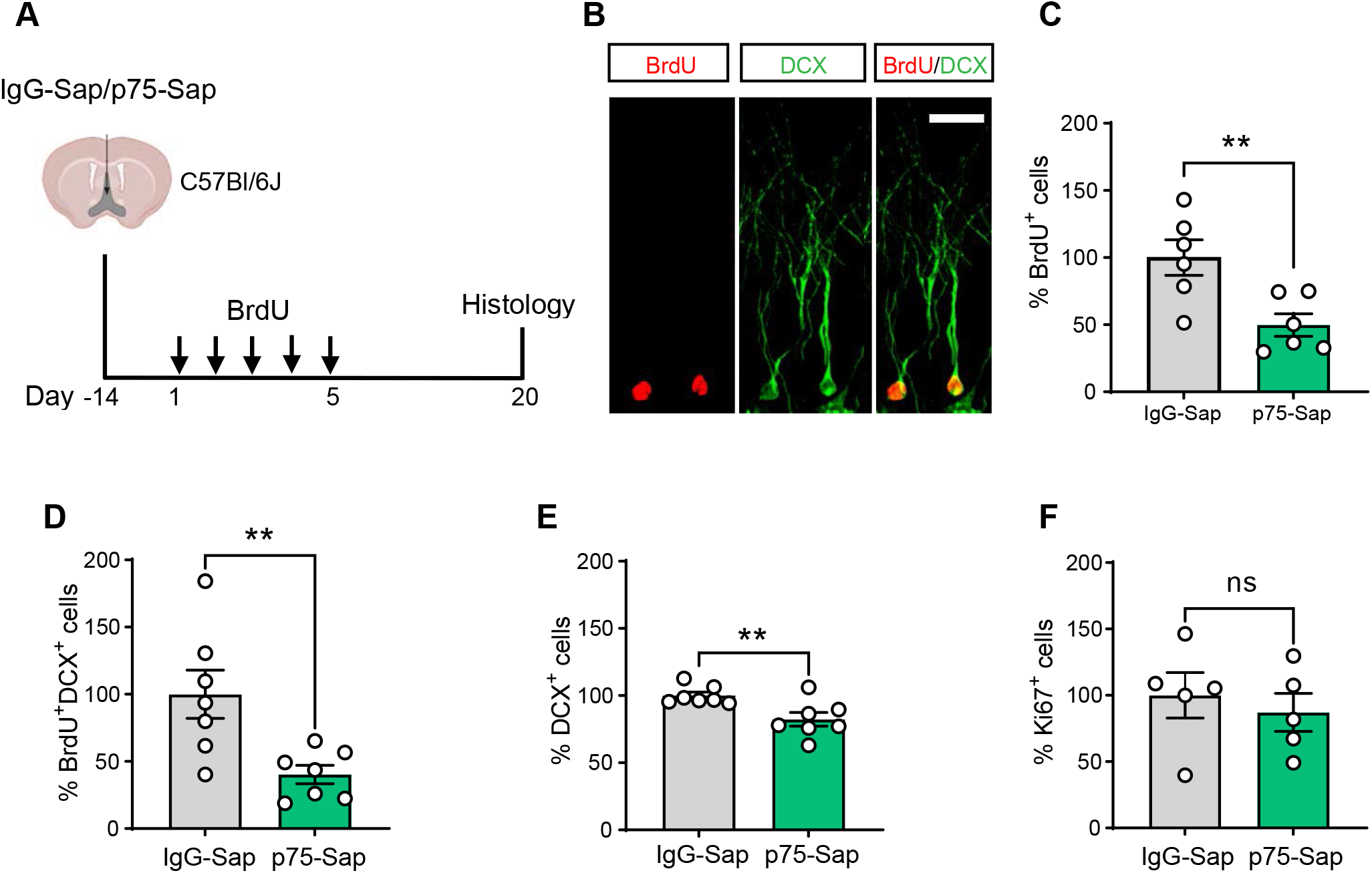
Selective MS/DBB cholinergic lesion leads to impairments in adult hippocampal neurogenesis. (A) Overview of the experimental timeline to evaluate adult neurogensis. Two weeks following IgG-Sap or p75-Sap infusion in the MS/DBB, mice received daily injections of BrdU for five days and were sacrificed three weeks later. (B) A confocal image showing newborn hippocampal neurons co-labeled with BrdU (red) and DCX (green). (C-E) Quantification represented as ‘percentage change’ relative to IgG-Sap-infused controls. Note the significant reduction in the total number of BrdU^+^ cells (C), newly generated immature neurons (BrdU^+^DCX^+^) (D) and total immature neurons (DCX^+^) (E) in p75-Sap-infused mice. (F) No difference in the total number of proliferating cells (Ki67^+^) was observed between the groups. Data represent mean ± SEM. \*\**p* < 0.01; unpaired Student’s t-test. ns, not significant. Scale bar: 5 µm.

### Loss of MS/DBB cholinergic neurons impairs the morphological maturation of newborn neurons in the hippocampus

Cholinergic signaling has been shown to play an important role in shaping postnatal neural development, including neuronal maturation and dendritic development (Picciotto et al., 2012). Therefore, we next investigated whether the loss of MS/DBB cholinergic projections alters the morphological maturation of the surviving newborn hippocampal neurons. To address this, we adopted a genetic strategy involving Ascl1^CreERT2^::tdTom mice to label and lineage trace a cohort of adult-born (tdTom^+^) neurons (Yang et al., 2015; Jhaveri et al., 2018). Two groups of Ascl1^CreERT2^::tdTom mice were set up, with one group receiving a p75-Sap and the other an IgG-Sap injection in the MS/DBB. To characterize the morphology of immature adult-born neurons, these mice were sacrificed 4 weeks following the administration of tamoxifen (Fig. 3A) and the dendrites of individual tdTom^+^ neurons were traced and analyzed (Fig. 3B, C). We observed a significant reduction in the total dendritic length and the length of the apical (longest) dendrite of tdTom^+^ newborn neurons in the p75-Sap-injected mice compared to the control mice (Fig. 3D, E). Furthermore, Sholl analysis revealed a significant difference in the dendritic arborization between groups, with tdTom^+^ neurons from the p75-Sap group exhibiting branching closer to the cell body than neurons from the IgG-Sap group (Fig. 3G). However, the total number of dendritic endings and intersections were similar between groups (Fig. 3F). These data demonstrate that MS/DBB cholinergic innervation is important for the morphological maturation of newborn hippocampal neurons, with its loss leading to impairments in their dendritic development and branching pattern.

**Figure 3.**
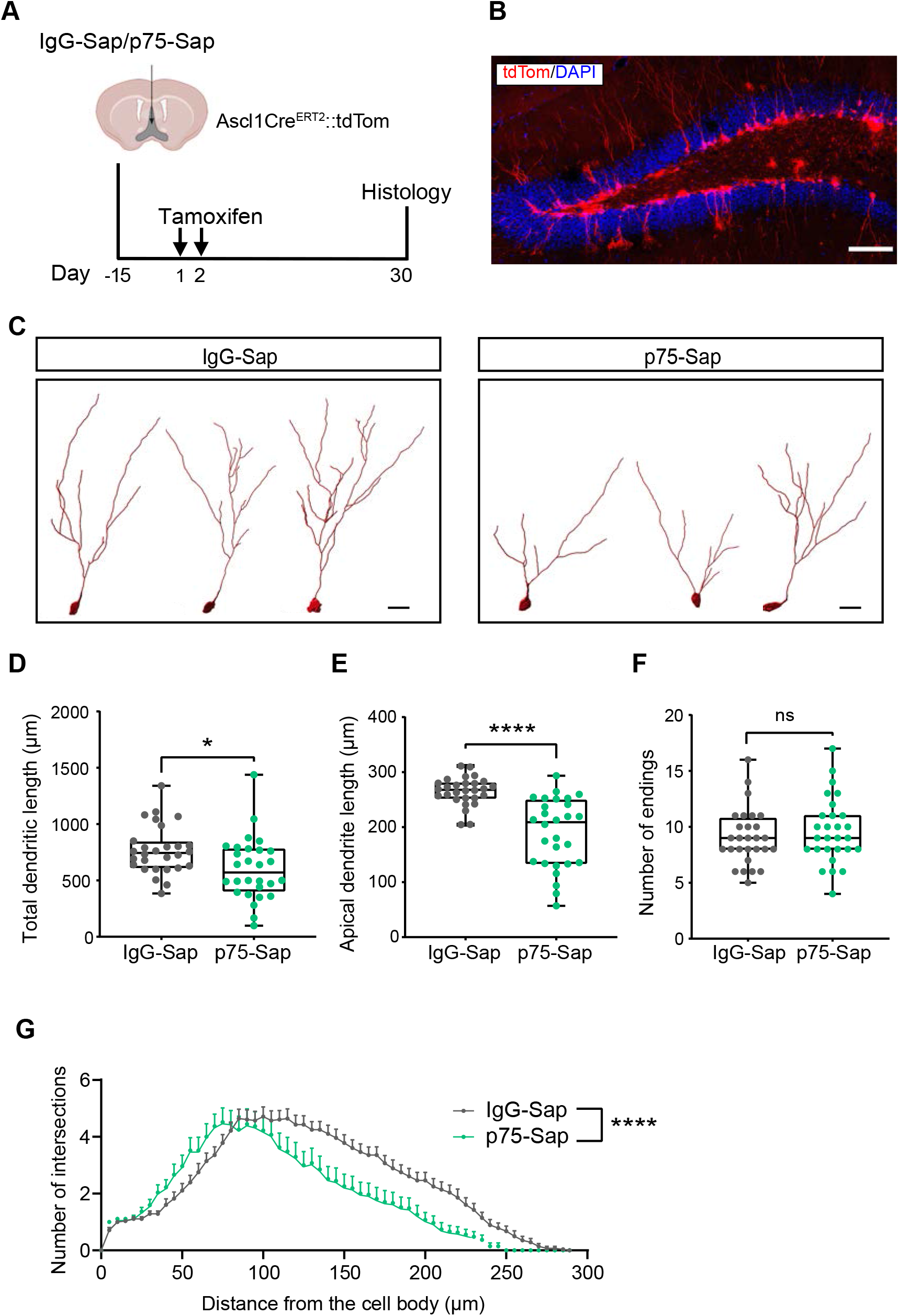
Loss of MS/DBB cholinergic neurons impairs dendritic development of immature adult-born neurons in the hippocampus. (A) Experimental timeline showing labeling of adult-born neurons using Ascl1^CreERT2^::tdTom mice. Two weeks after IgG-Sap or p75-Sap infusion in the MS/DBB, mice were injected with tamoxifen for two consecutive days to label and lineage trace new neurons. They were then sacrificed four weeks later. (B) A confocal image showing adult-born neurons (tdTom^+^, red) and nuclei stained with DAPI (blue) in the hippocampus. Scale bar: 100µm (C) Representative tracing of tdTom^+^ adult-born neurons from IgG-Sap and p75-Sap-treated mice. Scale bar: 20 μm (D) A significant reduction in the total dendritic length and (E) length of the apical dendrite was observed in the neurons of the p75-Sap-treated mice. (F) No difference in the number of endings of the immature adult-born neurons was noted between groups. (G) Sholl analysis revealed a significant difference in the dendritic architecture of up to 4-week-old adult-born neurons in p75-Sap-treated compared to control mice. n = 4 mice per group; 7 neurons per mouse; two-way ANOVA. Data represent mean ± SEM. \**p* < 0.05, \*\*\*\**p* < 0.0001; unpaired Student’s t-test (D-F).

### Pharmacological stimulation of Chrm4 directly activates a subpopulation of quiescent hippocampal NPCs *ex vivo*

Having established that the loss of cholinergic signaling from the MS/DBB impairs the process of adult hippocampal neurogenesis, resulting in fewer and morphologically altered newborn neurons, we next questioned whether this impairment could be ameliorated by directly modulating the activity of a cholinergic receptor subtype expressed on the hippocampal NPCs. We first stimulated hippocampal NPCs with drugs targeting the two major classes of acetylcholine receptors on these cells, namely metabotropic muscarinic and ionotropic nicotinic receptors, using the neurosphere assay (Fig. 4A). A single-cell suspension of primary hippocampal cells was treated with or without muscarine or nicotine in the presence of the growth factors EGF and bFGF (control condition). Treatment with 50 µM muscarine but not nicotine led to a significant increase in the number of neurospheres compared to the control condition, indicating that mAChRs may regulate the activity of hippocampal NPCs (Fig. 4B).

**Figure 4.**
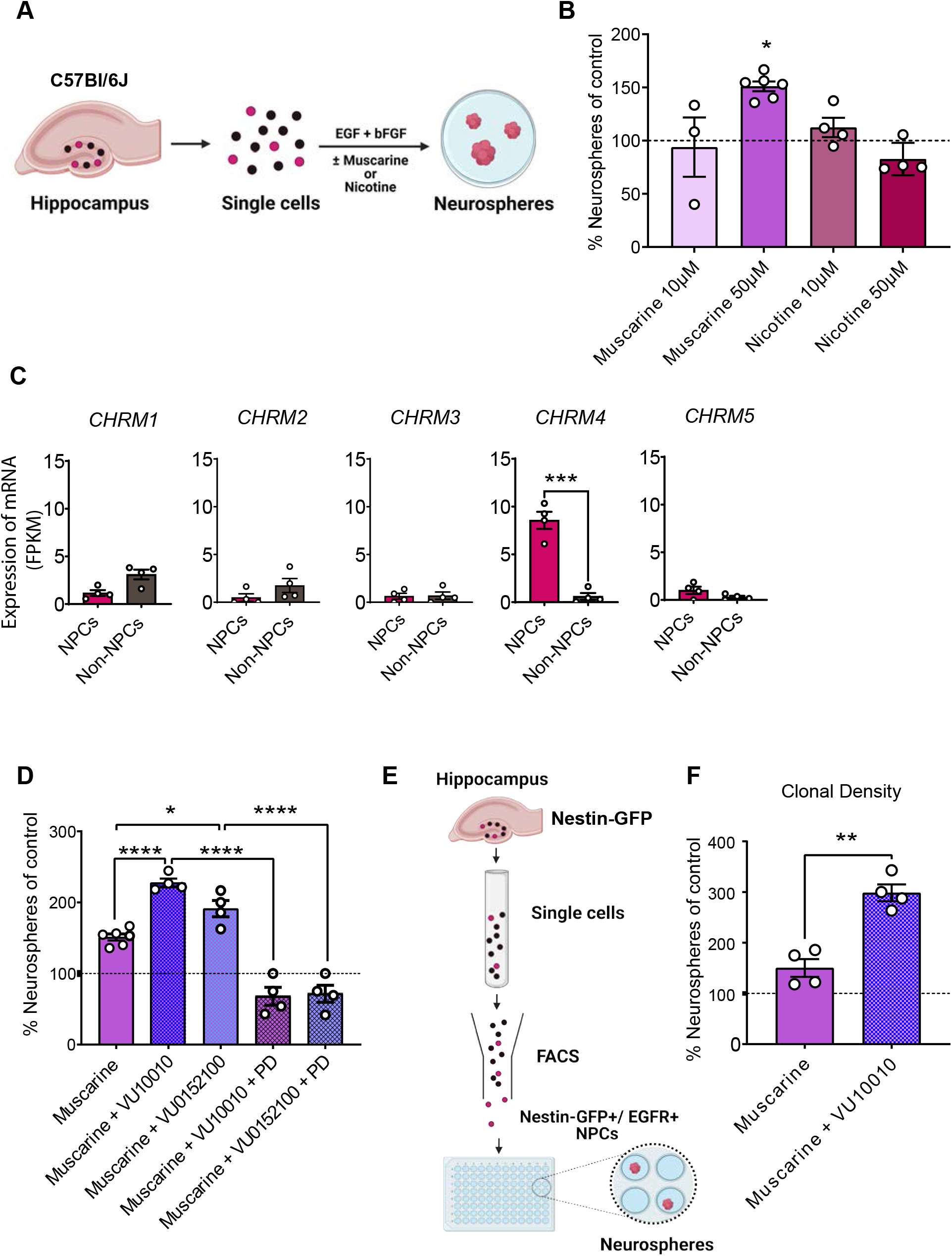
Selective stimulation of M4 mAChRs directly activates a population of quiescent hippocampal NPCs *ex vivo*. (A) Schematic of the neurosphere assay used to assess NPC activity from the hippocampus. (B) Treatment of adult hippocampal cells with muscarine (50 μM) but not nicotine generated significantly more neurospheres than produced in the control medium. (C) RNA-seq data showing expression of all muscarinic receptors (*CHRM1-5*) in the hippocampal NPC (Nestin-GFP^+^/EGFR^+^) and non-NPC (Nestin-GFP^−^/EGFR^−^) populations. Note the selective expression of *CHRM4* in the hippocampal NPCs. (paired Student’s t-test) (D) Treatment with the M4 mAChRs-selective allosteric potentiators VU10010 (10 μM) and VU0152100 (10 μM) in the presence of muscarine (50 μM) led to a two-fold increase in the number of neurospheres compared to the control (shown as dotted line). PD102807 (PD; 1 μM), a selective M4 mAChR antagonist, completed blocked this VU-mediated increase in neurosphere formation (one-way ANOVA). (E) Schematic showing the experimental design for the purification of Nestin-GFP^+^/EGFR^+^ NPCs from the adult hippocampus using FACS and the assay to assess their proliferative capacity at clonal density. (F) In the clonal density assay, the treatment of Nestin-GFP^+^/EGFR^+^ NPCs with VU10010 (10 μM) directly potentiated the effects of muscarine, with the relative number of neurospheres obtained in the presence of muscarine + VU10010 being significantly higher than that in muscarine alone (paired Student’s t-test). Data represent mean ± SEM. \**p* < .05, \*\**p* < 0.01, \*\*\**p* < 0.001, \*\*\**p* < 0.001.

We next sought to identify the cholinergic receptor(s) expressed by NPCs and examine the effects of selective stimulation of muscarinic receptor(s) on NPC activity. We have previously identified and purified both active and quiescent hippocampal NPCs using concomitant selection of Nestin-GFP^+^ and EGFR^+^ cells (Jhaveri et al., 2015). RNA-seq analysis of this population revealed selective expression and enrichment of *CHRM4* (the gene encoding M4 mAChR) but not *CHRM1, CHRM2, CHRM3* or *CHRM5* in Nestin-GFP^+^EGFR^+^ hippocampal NPCs compared to non-NPCs (Fig. 4C). Corroborating our data from the neurosphere assay, no or negligible expression of nicotine receptors was noted in this NPC population (data not shown).

Having identified that the M4 mAChR was preferentially expressed in the hippocampal NPCs, we then used pharmacological agents that selectively stimulate (VU10010 and VU0152100) or inhibit (PD102807) its activity in NPC cultures. As observed earlier, treatment of NPCs with muscarine alone led to a significant increase in the number of neurospheres, with the M4 mAChR-specific allosteric modulators VU10010 or VU0152100 further potentiating this effect (Fig. 4D). However, in the presence of PD102807, a selective M4 mAChR antagonist, this increase was completely abrogated, thereby establishing the specificity of the response (Fig. 4D). These findings suggest that cholinergic signaling via the M4 mAChR can activate a population of quiescent hippocampal NPCs.

Finally, to determine whether M4 mAChR stimulation exerts its effects directly or indirectly, hippocampal NPCs (Nestin-GFP^+^/EGFR^+^) were isolated using flow cytometry and plated at clonal density (Fig. 4E). At this density, treatment with VU10010 in the presence of muscarine still led to a significant increase in the number of neurospheres compared to the control (Fig. 4F). These findings indicate that cholinergic signaling via the M4 mAChR directly activates a population of hippocampal NPCs.

### Selective stimulation of the M4 mAChR enhances adult hippocampal neurogenesis *in vivo*

We next examined whether the stimulation of NPCs with selective M4 mAChR allosteric potentiators leads to an enhancement in the production of new neurons *in vivo*. VU10010, which is blood-brain barrier impermeable, or vehicle was infused directly into the hippocampus over a period of 7 days via a unilateral cannula (Fig. 5A). To label the proliferating cells, BrdU was administered daily for a period of 5 days, starting 48 h after the cannula implantation, and brains were collected 15 days after the end of infusion period. Subsequent analysis revealed a significant increase in the number of newly generated neurons (BrdU^+^DCX^+^) and the total number of immature neurons (DCX^+^) in the VU10010-compared to the vehicle-infused mice (Fig. 5B-D). This indicates that selective enhancement of M4 mAChR activity in the hippocampus enhances the production and/or survival of new neurons.

**Figure 5.**
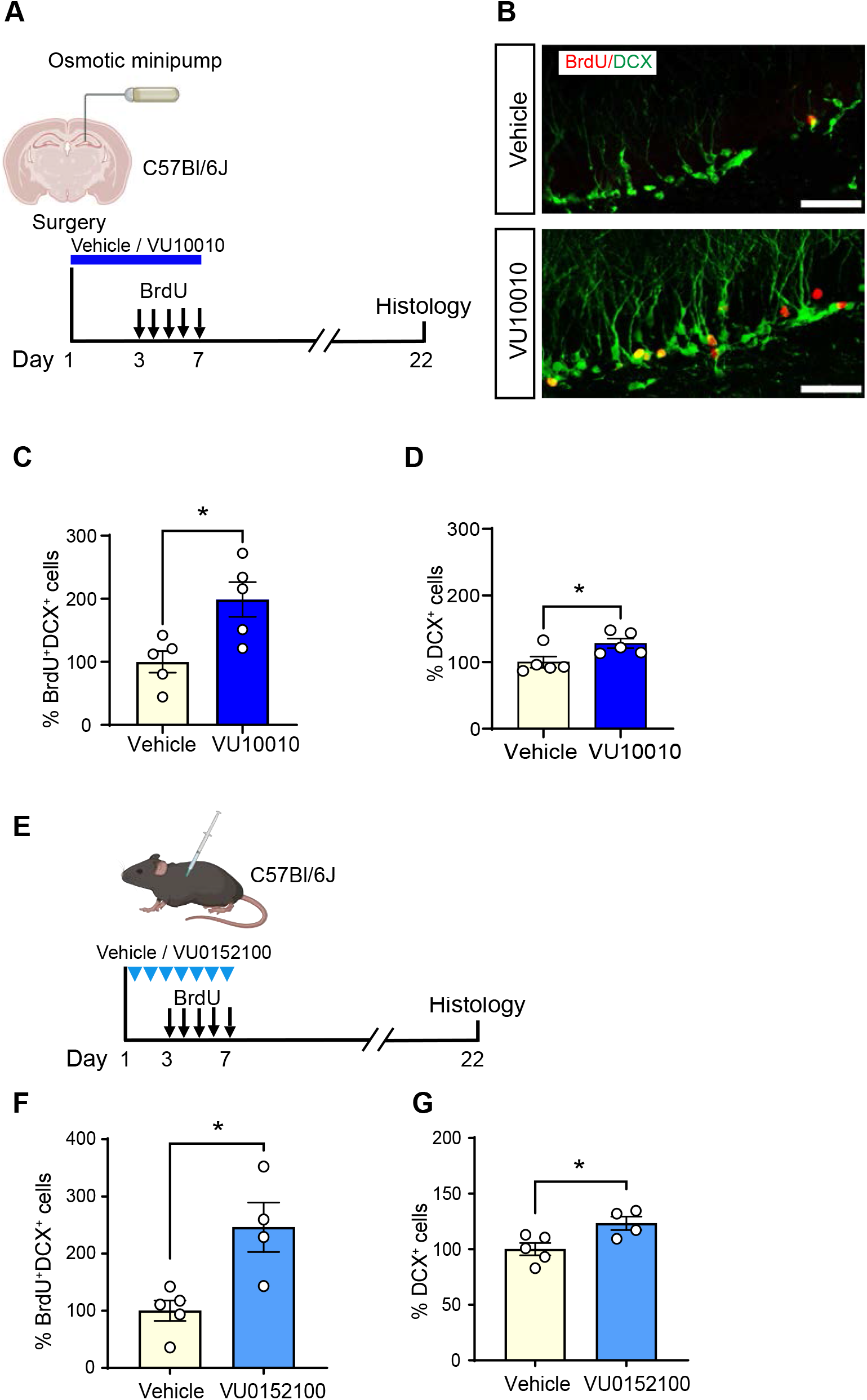
Selective stimulation of M4 mAChRs enhances adult hippocampal neurogenesis *in vivo*. (A) Experimental timeline for the unilateral infusion of VU10010 into the hippocampus. (B) Representative confocal images of the dentate gyrus showing BrdU^+^ (red) and DCX^+^ (green) cells, in the vehicle- and VU10010-treated mice. Scale bar: 50 µm. (C) Quantification showing a significant increase in the number of newborn neurons (BrdU^+^DCX^+^) and (D) total immature neurons (DCX^+^) in the VU10010-versus vehicle-infused hippocampus. (E) Experimental timeline for the systemic administration of VU0152100 to analyze the production of new neurons in the hippocampus. (F) The numbers of newborn neurons (BrdU^+^DCX^+^) and (G) total immature neurons (DCX^+^) were significantly higher in the VU0152100-treated mice. Data represent mean ± SEM: \**p* < 0.05; unpaired Student’s t-test.

To further corroborate these findings, we administered the blood-brain-barrier-permeable M4 mAChR potentiator VU0152100 (Brady et al., 2008) systemically via i.p. injections daily for 7 days (Fig. 5E). Similar to the results obtained following the infusion of VU10010, significant increases in the number of BrdU^+^DCX^+^ cells as well as the total number of DCX^+^ cells were observed in the VU0152100-treated mice compared to the controls (Fig. 5F, G).

Collectively, these data demonstrate that pharmacological stimulation of the M4 mAChR *in vivo* enhances the proliferation and/or survival of neurogenic NPCs and their progeny, boosting the overall level of adult hippocampal neurogenesis.

### Stimulating M4 mAChRs partially ameliorates MS/DBB cholinergic lesion-induced impairments in adult hippocampal neurogenesis

Given the pro-neurogenic effects of the M4 mAChR potentiators, we then investigated whether impairments in hippocampal neurogenesis observed following MS/DBB cholinergic neuron loss could be compensated by activating NPCs via selective stimulation of these mAChRs *in vivo*. We first selectively ablated MS/DBB cholinergic neurons by administering a single injection of p75-Sap. 14 days post-injection, mice were systemically administered either VU0152100 or vehicle daily for a period of 7 days (Fig. 6A). A third group of mice injected with IgG-Sap received vehicle injection and was used as the control. Similar to our earlier findings (Fig. 2), we observed a significant reduction in both the number of newborn neurons (BrdU^+^DCX^+^) and the total number of immature neurons (DCX^+^) in the p75-Sap+vehicle group compared to the IgG-Sap+vehicle group (Fig. 6B, C). However, we found no significant difference in either the number of BrdU^+^DCX^+^ newborn neurons or the total number of DCX^+^ immature neurons in p75-Sap+VU0152100-treated mice when compared to the control group (IgG-Sap+vehicle), suggesting that pharmacological stimulation of M4 mAChRs could compensate for the MS/DBB cholinergic neuron loss and restore the level of hippocampal neurogenesis to near basal levels.

**Figure 6.**
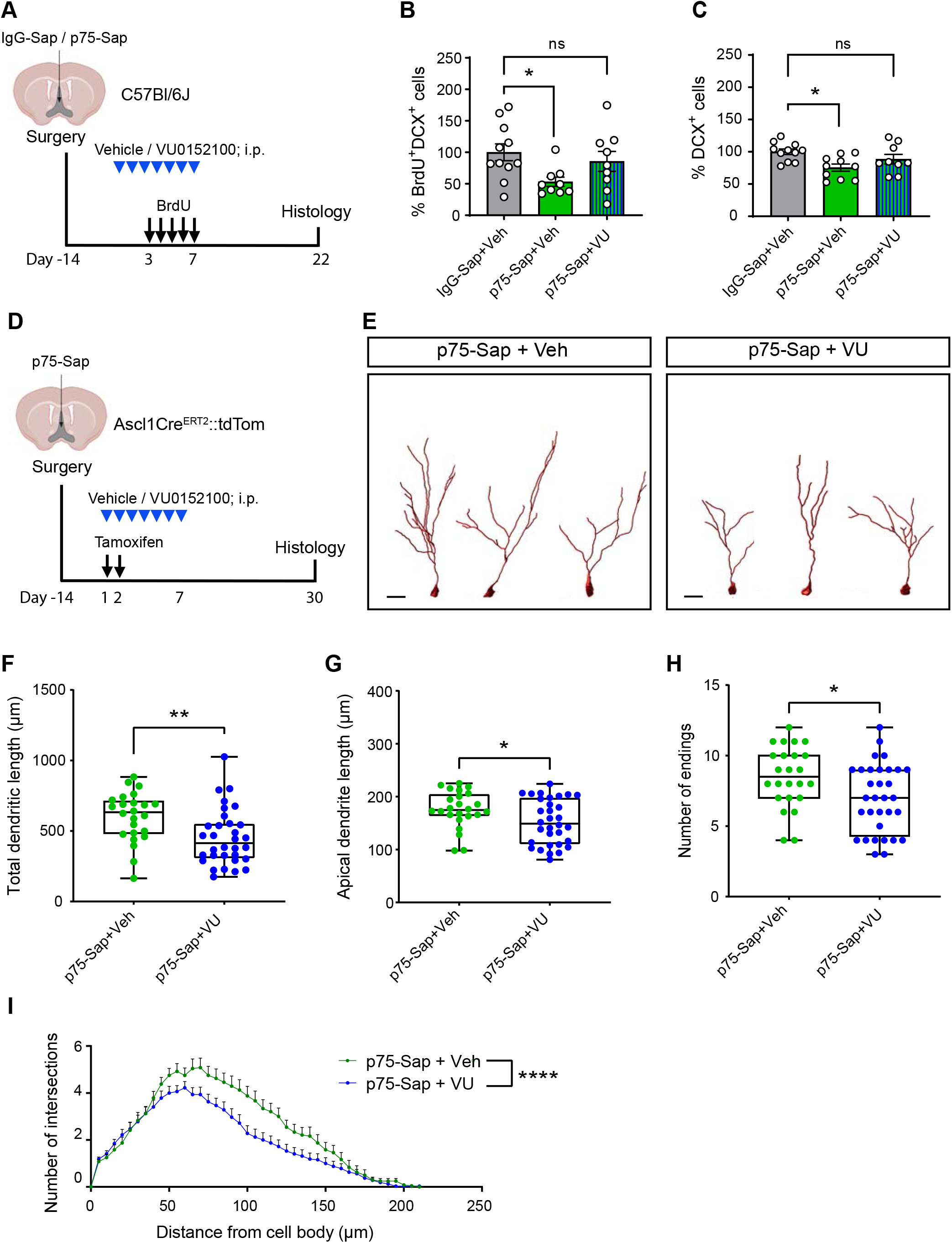
Effects of M4 mAChRs stimulation on adult hippocampal neurogenesis in MS/DBB cholinergic-lesioned mice. (A) Experimental timeline. 14 days post-IgG-Sap or p75-Sap infusion, mice were systemically administered either VU0152100 or vehicle daily for a period of 7 days. BrdU was administered daily for 5 consecutive days and mice were sacrificed 3 weeks later. (B) Quantification of the number of newborn neurons (BrdU^+^DCX^+^) and (C) total immature neurons (DCX^+^) in the vehicle- and VU015200-treated mice. Note the significant decrease in newly generated and total immature neurons in the p75-Sap+Veh-treated mice compared to the IgG-Sap+Veh-treated mice, whereas these numbers did not differ between IgG-Sap+Veh-treated and p75-Sap+VU-treated mice. (\**p* < 0.05; one-way ANOVA) (D) Experimental timeline for assessing the dendritic morphology of newborn neurons following MS/DBB cholinergic lesion and systemic administration of VU0152100 or vehicle using Ascl1^CreERT2^::tdTom mice. (E) Representative tracing of tdTom^+^ adult-born neurons from p75-Sap+Veh- and p75-Sap+VU-treated mice. Scale bar: 20 μm. (F) A significant reduction in the total dendritic length, (G) length of the apical dendrite and (H) number of endings of the immature neurons was found in p75-Sap+VU-compared to p75-Sap+Veh-treated mice. (I) Sholl analysis also revealed a significant decrease in the dendritic arborization of adult-born neurons in p75-Sap+VU-treated mice (n = 3 - 4 mice per group, 8 neurons per mouse; two-way ANOVA). Data represent mean ± SEM: \**p* <0.05, \*\**p* <0.01, \*\*\*\**p* <0.0001; unpaired Student’s t-test (F-H).

Finally, to examine whether selective M4 mAChR stimulation could also restore the appropriate morphological development of newborn neurons following MS/DBB cholinergic lesion, we used our Ascl1^CreERT2^::tdTom mice (Fig. 6D). Following p75-Sap-induced MS/DBB cholinergic neuron lesion, one group of mice received VU0152100 and the other vehicle for a period of 7 days. To assess the dendritic architecture of up to 4-week-old immature adult-born neurons, mice were sacrificed 4 weeks after tamoxifen administration. Strikingly, and in opposition to our hypothesis, we observed a significant reduction in the total dendritic length and the length of the apical dendrite of newborn neurons in the VU0152100-compared to the vehicle-treated mice (Fig. 6E-G). The total number of dendritic endings was also significantly reduced (Fig. 6H), with Sholl analysis revealing a significant decrease in the number of dendritic intersections in the VU0152100-treated mice (Fig. 6I).

Taken together, these findings suggest that, although selective stimulation of M4 mAChRs ameliorates the MS/DBB cholinergic lesion-induced decrease in the level of hippocampal neurogenesis, it does so without enhancing dendritic maturation. Rather, M4 mAChR stimulation further exacerbates the impairments in the morphological maturation of these adult-born neurons.

## Discussion

The production of new neurons in the dentate gyrus of the adult hippocampus is a multi-stage process that includes the activation and proliferation of quiescent NPCs within the neurogenic niche, their specification into neurons, and their functional differentiation and integration within the hippocampal circuit. This study has revealed the contributions of cholinergic signaling during at least three of these stages: (i) the selective expression of M4 mAChRs by quiescent NPCs, and the ability of M4 mAChRs-selective potentiators to activate these cells, resulting in increased numbers of newborn neurons; (ii) the necessity of direct septal-hippocampal cholinergic neurotransmission for the survival of adult-born neurons, and (iii) the ability of cholinergic stimulation (by direct innervation and through M4 mAChRs) to sculpt the dendritic architecture of newly generated immature neurons.

Our finding that the selective loss of MS/DBB cholinergic neurons in mice results in a significant impairment in adult hippocampal neurogenesis reflects that previously reported in rats following a global basal forebrain cholinergic lesion (Cooper-Kuhn et al., 2004; Mohapel et al., 2005). This demonstrates that the cholinergic inputs into the hippocampus from the MS/DBB directly regulate neurogenesis and shows that the regulation is not due to an indirect effect of the NBM or SI cholinergic neurons, although this remains a possible additional regulatory mechanism. Whereas our targeted approach resulted in a profound loss of septal-hippocampal innervation, a previous study that achieved only partial elimination of the MS/DBB cholinergic neurons was achieved reported no effect on baseline hippocampal neurogenesis (Ho et al., 2009). This indicates that a normal to low level of cholinergic innervation is sufficient to support baseline levels of hippocampal neurogenesis. Hence, strategies aimed at promoting the survival of MS/DBB cholinergic neurons, or mimicking the key lost cholinergic functions, could prove beneficial for the maintenance of ongoing neurogenesis and neurogenesis-dependent functions in various neurodegenerative conditions, including Alzheimer’s disease, in which basal forebrain cholinergic neuron loss is one of the key early pathogenic events (Hampel et al., 2019).

Our study revealed that NPCs selectively express M4 mAChRs with low to negligible expression of other cholinergic (muscarinic and nicotinic) receptors. Furthermore, pharmacological stimulation using M4-selective allosteric modulators (VU10010 and VU0152100) led to the activation of quiescent NPCs, resulting in increased production of new neurons in the adult hippocampus. Fibers expressing choline acetyltransferase, the enzyme responsible for acetylcholine synthesis, have been observed in close proximity to NPCs in the dentate gyrus (Kaneko et al., 2006), suggesting that these cholinergic responsive cells may receive direct inputs from the basal forebrain cholinergic neurons. M4 mAChRs are preferentially coupled to inhibitory G (G_i/o_) proteins, which can link with potassium channels to reduce the resting membrane potential (Ishii and Kurachi, 2006), as well as modulating several signaling pathways linked to altered rates of neurogenesis. For example, the membrane hyperpolarization of ventricular zone NPCs shifts the transcriptional program and neurogenic division mode (Vitali et al., 2018). Conversely, G_i/o_ proteins inhibit adenylate cyclase, leading to a decrease in intracellular cAMP (Langmead et al., 2008), which has been reported to act as a positive regulator of adult hippocampal neurogenesis through increased proliferation (Nakagawa et al., 2002). M4 mAChRs have also been shown to activate the mitogen-activated protein (MAP) kinase pathway, as well as transactivating the EGF receptor (Stirnweiss et al., 2006; Ockenga et al., 2013). Further studies will be required to elucidate whether any of these signaling mechanisms underpin the M4 mAChRs-mediated activation of quiescent hippocampal NPCs. Nonetheless, the current results provide the first demonstration of cholinergic-responsive quiescent NPCs in a neurogenic niche which, together with our previous identification of norepineprine- and dopamine-responsive subpopulations of quiescent hippocampal NPCs (Jhaveri et al., 2015), adds to the growing appreciation of neurotransmitter-mediated control of NPC activity in the adult hippocampus (Berg et al., 2013; Song et al., 2016).

We found that the loss of MS/DBB cholinergic innervation in the hippocampus led to a significant decrease in the number of newly generated neurons (BrdU^+^/DCX^+^) without altering the overall proliferation in the dentate gyrus, suggesting reduced survival of newly generated progenitors and/or neurons. Importantly, treatment with M4 mAChRs-selective allosteric potentiators rescued this phenotype, restoring the rates of adult neurogenesis in MS/DBB-lesioned mice. Moreover, both systemic administration (VU0152100) and intrahippocampal infusion (VU10010) of M4 modulators resulted in enhanced neurogenesis *in vivo* in non-lesioned animals. This could reflect the fact that stimulating quiescent M4 mAChRs-responsive NPCs leads to a neurogenic division mode (Vitali et al., 2018).

Our study also revealed that MS/DBB cholinergic neurons are important for the morphological maturation of surviving adult-born neurons, contributing to their structural plasticity. These findings add to the existing literature reporting the contribution of GABA and glutamate in the regulation of the dendritic development of adult-born neurons (Tashiro et al., 2006; Song et al., 2013), highlighting a major role of neurotransmitters in sculpting the morphological and connectivity patterns of these neurons in the hippocampus. In particular, results from the systemic administration of VU0152100 revealed that the stimulation of M4 mAChRs expressed by NPCs or their progeny exacerbated the reduced dendritic branching induced by loss of cholinergic innervation to the dentate gyrus. Given the inhibitory effect of M4 mAChRs on the neurotransmission of neurons, it is perhaps not surprising that the maturation and integration of newborn neurons is directly impaired. Interestingly, both loss of cholinergic function and activation of M2 and M4 mAChRs can suppress the hippocampal theta (or ripple) oscillations associated with hippocampal- and adult neurogenesis-dependent learning and memory (Berdugo-Vega et al., 2020; Kumar et al., 2020; Ma et al., 2020).

However, determining how septal cholinergic neurons regulate the morphological development of adult-born neurons in the hippocampus remains challenging, as this function likely involves both direct and indirect mechanisms. Retrograde tracing experiments using pseudo-type rabies virus have demonstrated the presence of significant and stable monosynaptic cholinergic inputs arising from the MS/DBB onto immature adult-born hippocampal neurons starting as early as 10 days after their birth (Deshpande et al., 2013). An indirect contribution of the basal forebrain cholinergic system to the modulation of adult neurogenesis via the entorhinal cortex has also been proposed (Mesulam et al., 1983). As robust glutamatergic synaptic inputs from the entorhinal cortex onto adult-born neurons are established at a more mature stage, around 3 weeks after their birth (Vivar et al., 2012; Deshpande et al., 2013), the indirect contribution of cholinergic signaling via these afferents may well regulate the later stages of maturation and/or integration of these neurons. Adding another layer of complexity is the finding that cholinergic terminals in the hippocampus co-transmit acetylcholine and GABA (Takács et al., 2018), raising the possibility that the impairments in the morphological development of adult-born neurons observed following MS/DBB cholinergic lesion could also be due to a reduction in GABAergic signaling. In support of this notion, GABAergic synaptic inputs from local parvalbumin-expressing interneurons have previously been shown to promote the survival and morphological development of adult-born neurons in the hippocampus (Song et al., 2013). Therefore, future studies that examine the expression of cholinergic receptor subtype(s) on immature adult-born neurons and determine their functional roles will be essential for uncovering the contribution of cholinergic signaling to the regulation of the structural plasticity of these cells.

Based on our findings, we propose that the neurogenesis-promoting activity of M4-selective allosteric potentiators might be of therapeutic benefit in conditions where reduced rates of neurogenesis are observed and neurogenesis-dependent functions are impaired such as during ageing and in neurodegenerative conditions. In fact, these M4 potentiators have been proposed as promising novel small molecule pharmaceuticals for the treatment of the cognitive and behavioral impairments present in Alzheimer’s disease and schizophrenia (Foster and Conn, 2017). However, our data showing excerbated impairments in the dendritic architecture of adult-born neurons following systemic administration of VU0152100 to MS/DBB-lesioned animals highlights the need to proceed with caution, especially when considering therapeutic strategies for conditions which are associated with the loss of basal forebrain cholinergic neurons.

In summary, our findings show that the MS/DBB cholinergic neurons which innervate the hippocampus play a crucial role in the stage-specific regulation of the neurogenic process, particularly supporting the survival and morphological maturation of newly generated neurons. In addition, we have uncovered cholinergic-responsive quiescent NPCs that can be activated via M4 mAChR-selective stimulation to enhance the generation of new neurons. Although these M4 mAChR-expressing quiescent NPCs could be recruited to ameliorate the MS/DBB cholinergic lesion-induced decrease in the levels of hippocampal neurogenesis, our findings uncouple the role of M4 mAChR-selective potentiators in regulating NPC activity from that which supports the morphological maturation of newborn neurons, at least in conditions where there is cholinergic neuron loss. We therefore propose that future studies not only examine the roles of select compounds/genes/signaling in altering the overall level of neurogenesis but also extend their investigation to include an assessment of the structural and functional properties of adult-born neurons to obtain a comprehensive understanding of the stage-specific regulation of neurogenesis and its implications for the modulation of hippocampus-dependent functions.

## Acknowledgments

This work was supported by a Mater Foundation Fellowship and Dementia Australia Research Foundation Project Grant to D.J.J. We thank the Queensland Brain Institute (QBI) Animal Facility staff for breeding and maintaining the animals used in this study and acknowledge the QBI and Translational Research Institute Flow Cytometry Facilities for technical assistance. Imaging was performed at the QBI’s Advanced Microscopy Facility, generously supported by the Australian Government through the ARC LIEF grant LE100100074. We are grateful to Rowan Tweedale for editorial assistance.

## Author contributions

L.I.M., E.J.C. and D.J.J. designed the project; L.I.M., S.B., K.H., J.J.M., M.M. and D.J.J. performed experiments and analysed data; L.I.M. and D.J.J. prepared the original draft; E.J.C. and D.J.J. reviewed and edited the manuscript and provided supervision; D.J.J. obtained funding.

## References

Auld DS, Kornecook TJ, Bastianetto S, Quirion R (2002) Alzheimer’s disease and the basal forebrain cholinergic system: relations to β-amyloid peptides, cognition, and treatment strategies. Progress in Neurobiology 68:209–245.

Berdugo-Vega G, Arias-Gil G, López-Fernández A, Artegiani B, Wasielewska JM, Lee C-C, Lippert MT, Kempermann G, Takagaki K, Calegari F (2020) Increasing neurogenesis refines hippocampal activity rejuvenating navigational learning strategies and contextual memory throughout life. Nat Commun 11:135.

Berg DA, Belnoue L, Song H, Simon A (2013) Neurotransmitter-mediated control of neurogenesis in the adult vertebrate brain. Development 140:2548–2561.

Brady AE, Jones CK, Bridges TM, Kennedy JP, Thompson AD, Heiman JU, Breininger ML, Gentry PR, Yin H, Jadhav SB, Shirey JK, Conn PJ, Lindsley CW (2008) Centrally active allosteric potentiators of the M4 muscarinic acetylcholine receptor reverse amphetamine-induced hyperlocomotor activity in rats. J Pharmacol Exp Ther 327:941–953.

Chen BH, Ahn JH, Park JH, Choi SY, Lee YL, Kang IJ, Hwang IK, Lee TK, Shin BN, Lee JC, Hong S, Jeon YH, Shin MC, Cho JH, Won MH, Lee YJ (2018) Effects of Scopolamine and Melatonin cotreatment on cognition, neuronal damage, and neurogenesis in the mouse dentate gyrus. Neurochem Res 43:600–608.

Christian KM, Song H, Ming GL (2014) Functions and dysfunctions of adult hippocampal neurogenesis. Annu Rev Neurosci 37:243–262.

Cooper-Kuhn CM, Winkler J, Kuhn HG (2004) Decreased neurogenesis after cholinergic forebrain lesion in the adult rat. Journal of neuroscience research 77:155–165.

Deshpande A, Bergami M, Ghanem A, Conzelmann K-K, Lepier A, Götz M, Berninger B (2013) Retrograde monosynaptic tracing reveals the temporal evolution of inputs onto new neurons in the adult dentate gyrus and olfactory bulb. Proceedings of the National Academy of Sciences 110:E1152–E1161.

Foster DJ, Conn PJ (2017) Allosteric modulation of GPCRs: New insights and potential utility for treatment of schizophrenia and other CNS disorders. Neuron 94:431–446.

Hamlin AS, Windels F, Boskovic Z, Sah P, Coulson EJ (2013) Lesions of the basal borebrain Cholinergic System in Mice Disrupt Idiothetic Navigation. PLoS One 8:e53472.

Hampel H, Mesulam MM, Cuello AC, Khachaturian AS, Vergallo A, Farlow MR, Snyder PJ, Giacobini E, Khachaturian ZS (2019) Revisiting the cholinergic hypothesis in Alzheimer’s Disease: emerging evidence from translational and clinical research. Journal of Prevention of Alzheimer’s Disease 6:2–15.

Ho NF, Han SP, Dawe GS (2009) Effect of voluntary running on adult hippocampal neurogenesis in cholinergic lesioned mice. BMC Neuroscience 10:57.

Ishii M, Kurachi Y (2006) Muscarinic acetylcholine receptors. Curr Pharm Des 12:3573–3581.

Jhaveri DJ, Mackay EW, Hamlin AS, Marathe SV, Nandam LS, Vaidya VA, Bartlett PF (2010) Norepinephrine directly activates adult hippocampal precursors via beta3-adrenergic receptors. J Neurosci 30:2795–2806.

Jhaveri DJ, Tedoldi A, Hunt S, Sullivan R, Watts NR, Power JM, Bartlett PF, Sah P (2018) Evidence for newly generated interneurons in the basolateral amygdala of adult mice. Mol Psychiatry 23:521–532.

Jhaveri DJ, O’Keeffe I, Robinson GJ, Zhao QY, Zhang ZH, Nink V, Narayanan RK, Osborne GW, Wray NR, Bartlett PF (2015) Purification of neural precursor cells reveals the presence of distinct, stimulus-specific subpopulations of quiescent precursors in the adult mouse hippocampus. J Neurosci 35:8132–8144.

Jhaveri DJ, Taylor C.J., Bartlett P.F. (2012) Activation of different neural precursor populations in the adult hippocampus: does this lead to new neurons with discrete functions? Dev Neurobiol 72(7), 1044–1058.

Kaneko N, Okano H, Sawamoto K (2006) Role of the cholinergic system in regulating survival of newborn neurons in the adult mouse dentate gyrus and olfactory bulb. Genes to cells : devoted to molecular & cellular mechanisms 11:1145–1159.

Kerbler GM, Hamlin AS, Pannek K, Kurniawan ND, Keller MD, Rose SE, Coulson EJ (2013) Diffusion-weighted magnetic resonance imaging detection of basal forebrain cholinergic degeneration in a mouse model. Neuroimage 66:133–141.

Kotani S, Yamauchi T, Teramoto T, Ogura H (2006) Pharmacological evidence of cholinergic involvement in adult hippocampal neurogenesis in rats. Neuroscience 142:505–514.

Kumar D et al. (2020) Sparse activity of hippocampal adult-born neurons during REM sleep is necessary for memory consolidation. Neuron 107:552-565.e510.

Langmead CJ, Watson J, Reavill C (2008) Muscarinic acetylcholine receptors as CNS drug targets. Pharmacology & Therapeutics 117:232–243.

Lugert S, Basak O, Knuckles P, Haussler U, Fabel K, Gotz M, Haas CA, Kempermann G, Taylor V, Giachino C (2010) Quiescent and active hippocampal neural stem cells with distinct morphologies respond selectively to physiological and pathological stimuli and aging. Cell Stem Cell 6:445–456.

Ma X, Zhang Y, Wang L, Li N, Barkai E, Zhang X, Lin L, Xu J (2020) The firing of theta state-related septal cholinergic neurons disrupt hippocampal ripple oscillations via muscarinic receptors. J Neurosci 40:3591–3603.

Madrid LI, Jimenez-Martin J, Coulson EJ, Jhaveri DJ (2021) Cholinergic regulation of adult hippocampal neurogenesis and hippocampus-dependent functions. Int J Biochem Cell Biol 134:105969.

Mesulam MM, Mufson EJ, Wainer BH, Levey AI (1983) Central cholinergic pathways in the rat: an overview based on an alternative nomenclature (Ch1-Ch6). Neuroscience 10:1185–1201.

Ming GL, Song H (2011) Adult neurogenesis in the mammalian brain: significant answers and significant questions. Neuron 70:687–702.

Mohapel P, Leanza G, Kokaia M, Lindvall O (2005) Forebrain acetylcholine regulates adult hippocampal neurogenesis and learning. Neurobiology of aging 26:939–946.

Moreau PH, Cosquer B, Jeltsch H, Cassel JC, Mathis C (2008) Neuroanatomical and behavioral effects of a novel version of the cholinergic immunotoxin mu p75-saporin in mice. Hippocampus 18:610–622.

Moreno-Jimenez EP, Flor-Garcia M, Terreros-Roncal J, Rabano A, Cafini F, Pallas-Bazarra N, Avila J, Llorens-Martin M (2019) Adult hippocampal neurogenesis is abundant in neurologically healthy subjects and drops sharply in patients with Alzheimer’s disease. Nat Med 25:554–560.

Nakagawa S, Kim JE, Lee R, Malberg JE, Chen J, Steffen C, Zhang YJ, Nestler EJ, Duman RS (2002) Regulation of neurogenesis in adult mouse hippocampus by cAMP and the cAMP response element-binding protein. J Neurosci 22:3673–3682.

Ockenga W, Kühne S, Bocksberger S, Banning A, Tikkanen R (2013) Non-Neuronal Functions of the M2 Muscarinic Acetylcholine Receptor. Genes 4:171–197.

Picciotto MR, Higley MJ, Mineur YS (2012) Acetylcholine as a neuromodulator: cholinergic signaling shapes nervous system function and behavior. Neuron 76:116–129.

Schmitz TW, Nathan Spreng R (2016) Basal forebrain degeneration precedes and predicts the cortical spread of Alzheimer’s pathology. Nat Commun 7:13249.

Shirey JK, Xiang Z, Orton D, Brady AE, Johnson KA, Williams R, Ayala JE, Rodriguez AL, Wess J, Weaver D, Niswender CM, Conn PJ (2008) An allosteric potentiator of M4 mAChR modulates hippocampal synaptic transmission. Nat Chem Biol 4:42–50.

Song J, Olsen RH, Sun J, Ming GL, Song H (2016) Neuronal Circuitry Mechanisms Regulating Adult Mammalian Neurogenesis. Cold Spring Harb Perspect Biol 8.

Song J, Sun J, Moss J, Wen Z, Sun GJ, Hsu D, Zhong C, Davoudi H, Christian KM, Toni N, Ming G-l, Song H (2013) Parvalbumin interneurons mediate neuronal circuitry–neurogenesis coupling in the adult hippocampus. Nature Neuroscience 16:1728–1730.

Song J, Zhong C, Bonaguidi MA, Sun GJ, Hsu D, Gu Y, Meletis K, Huang ZJ, Ge S, Enikolopov G, Deisseroth K, Luscher B, Christian KM, Ming GL, Song H (2012) Neuronal circuitry mechanism regulating adult quiescent neural stem-cell fate decision. Nature 489:150–154.

Stirnweiss J, Valkova C, Ziesché E, Drube S, Liebmann C (2006) Muscarinic M2 receptors mediate transactivation of EGF receptor through Fyn kinase and without matrix metalloproteases. Cellular Signalling 18:1338–1349.

Takács VT, Cserép C, Schlingloff D, Pósfai B, Szőnyi A, Sos KE, Környei Z, Dénes Á, Gulyás AI, Freund TF, Nyiri G (2018) Co-transmission of acetylcholine and GABA regulates hippocampal states. Nat Commun 9:2848.

Tashiro A, Sandler VM, Toni N, Zhao C, Gage FH (2006) NMDA-receptor-mediated, cell-specific integration of new neurons in adult dentate gyrus. Nature 442:929–933.

Van Kampen JM, Eckman CB (2010) Agonist-induced restoration of hippocampal neurogenesis and cognitive improvement in a model of cholinergic denervation. Neuropharmacology 58:921–929.

Vitali I, Fièvre S, Telley L, Oberst P, Bariselli S, Frangeul L, Baumann N, McMahon JJ, Klingler E, Bocchi R, Kiss JZ, Bellone C, Silver DL, Jabaudon D (2018) Progenitor hyperpolarization regulates the sequential generation of neuronal subtypes in the developing neocortex. Cell 174:1264-1276.e1215.

Vivar C, Potter MC, Choi J, Lee J-Y, Stringer TP, Callaway EM, Gage FH, Suh H, van Praag H (2012) Monosynaptic inputs to new neurons in the dentate gyrus. Nat Commun 3:1107–1107.

Walker TL, White A, Black DM, Wallace RH, Sah P, Bartlett PF (2008) Latent stem and progenitor cells in the hippocampus are activated by neural excitation. J Neurosci 28:5240–5247.

Woolf NJ, Gould E, Butcher LL (1989) Nerve growth factor receptor is associated with cholinergic neurons of the basal forebrain but not the pontomesencephalon. Neuroscience 30:143–152.

Yang SM, Alvarez DD, Schinder AF (2015) Reliable genetic labeling of adult-born dentate granule cells using Ascl1 CreERT2 and Glast CreERT2 murine lines. J Neurosci 35:15379–15390.

